## Supplementary Note S1: Supplemental text, methods, and Supplemental Figures S1 - S16. for "An evolutionary approach to predict the orientation of CRISPR arrays"

### Supplementary Note 1:

**A. Simulation datasets & used parameters.** We follow the simulation process described in [1], Supplementary Note 1, closely.

We simulated two datasets to evaluate the quality of *CRISPR-evOr*:

**Dataset A: Simulations on coalescent trees.** A dataset similar to the one used in [1], which they used to evaluate the parameter estimation performance. Every group is composed of $k = 16$ samples and 500 trees were simulated according to the coalescent (with branch lengths exponentially distributed with mean $2/(k \cdot (k-1))$. Then we sampled the stationary length distribution of the block deletion model with ordered insertions (a Poisson distribution with mean $\theta/(\rho_\mathrm{B}\alpha)$, where $\theta$ is the insertion rate, $\rho_\mathrm{B}$ the block deletion initiation rate and $\alpha$ the average block length). Subsequently, we evolved this progenitor along the tree according to the block deletion model with ordered insertions as is described in [1], Supplementary Note 1. We chose the same parameters, i.e. insertion rate $\theta = 6$, average array length $n = 22$, $\alpha \in \{1, 1.1, \ldots, 4.9, 5\}$. The deletion initiation rate $\rho_\mathrm{B}$ was chosen such that the average per spacer deletion rate $\rho_\mathrm{B} \cdot \alpha$ remains constant ($\rho_\mathrm{B} = \theta/(n\alpha) = \text{const.}$). Then simulations were run for each of the 500 trees and each value of $\alpha$ for a total of 20500 simulated groups.

**Dataset B: Simulations on core-genome trees with estimated parameters.** As a base for this database, we relied upon the CRISPRCasdb dataset used in [1], composed of all groups (334) with at least one reconstructed deletion in the reconstruction, which is a subset of the dataset of 518 groups used for the evaluation of *CRISPR-evOr* in the main manuscript. We used the core-genome trees, estimated for each of these groups in [1], as underlying trees. Then we simulated as in A with parameters chosen according to the estimates for the respective group (with insertion rate estimated (crudely) by $\theta \approx n\rho_\mathrm{B}\alpha$ as described in [1]). We excluded a few groups with extreme parameter values and unstable simulations. For each group we ran 50 simulations and obtained a total of 13890 simulated groups.

As is described in the main manuscript, dataset A is used for a first evaluation of *CRISPR-evOr* where we obtained 100% accuracy for the orientation prediction. Dataset B was used for the rest of the simulation based evaluation, i.e. (a) runs of *CRISPR-evOr* using the original core-genome trees (97.8% accuracy) and (b) runs of *CRISPR-evOr* using SpacerPlacer to estimate trees (for each respective array orientation, 95.5% accuracy).

**B. Benchmarking on simulation datasets with conservative choice of decision threshold $c = 5$.** To round off the simulation experiments reported in the main manuscript (section "simulation-based benchmarking") we report the results for the conservative choice decision threshold of $c = 5$ (which was chosen according to manual investigation on the CRISPRCasdb dataset).

We get the following results for the three experiments using a threshold $c = 5$:

On dataset A, the lowest likelihood ratio $L_\text{ratio} \approx 26$ far exceeds the threshold $c = 5$, and thus there are no "not determined" or wrong predictions.

On dataset B (see above) using the original core-genome trees during inference of the orientation, we found that 95.5% of the predictions are correct and 4.5% are "not determined" and no wrong predictions are given.

On dataset B (see above) estimating the tree with SpacerPlacer during inference of the orientation, 91.5% of groups are predicted correctly, while 2.2% have the wrong orientation prediction and 6.3% are uncertain, i.e. "not determined".

Thus, comparing these results to the results in the main manuscript for $c = 0$, we are able to reduce the number of wrong predictions by *CRISPR-evOr* with conservative choices of $c$, with just a small trade-off in additional uncertain, i.e. "not determined", predictions.

**C. Remarks about the datasets used for group based count plots.** All orientation methods, excluding our own, consider only single arrays for their predictions. When considering groups (instead of individual arrays), not all arrays have a (confident) orientation prediction. However, we found no inconsistent predictions within groups for CRISPRstrand and PAM-orientation and only 3 groups with disagreeing orientation predictions for CRISPRDirection. Thus we identify the orientation of a group of arrays with the orientation of confident members of the group and extend the amount of available groups with orientation predictions. In cases of disagreement, we assign the orientation by majority decision. A jitterplot showing the fractions of confident predictions for each group and tool are shown in Supplementary Fig. S16.

Through this assignment, we increase the amount of arrays for which we are able to make orientation decisions substantially. For example, for the PAM dataset, this extends the dataset to the 77.2% group overlap (400 groups) that cover 81% of the arrays in the CRISPRCasdb dataset (instead of only considering the actual overlap of 38.9% of the arrays). Note, that this extension is only used for the group count plots and not for any of the array count based ones (here and in the main manuscript). Furthermore, this treatment of group predictions is favorable for the single array based methods (although grouping itself might not be).

**D. Remarks about predictions in csv and github examples.** All the reconstruction examples shown in the main manuscript or the supplement are part of the example datasets in the github repository ("orientation_paper_examples"). The example shown in Figure 2 in the main manuscript differs in the SpacerPlacer github repository, since we dropped one sample for better readability (LR890464).

Moreover, all of the examples are also available in the Supplementary csv with all orientation predictions. Namely, Figure 2 is g_768, Supplementary Fig. S2a is g_14, b is g_104 and S3 is g_848.
Note that the consensus repeats shown in the supplemental csv file of our results can be the reverse complements of the same samples in CRISPRCasdb. This can happen due to the clustering steps we used to arrive at our final datasets. We checked if there is overlap between groups with same consensus and reverse consensus repeat. If such groups were combined, we adjusted the orientation predictions for the arrays that were reversed to build the larger group.

### E. Description of columns in Table S1 csv file.

**group_name:** identifier of a group of arrays

**consensus_repeat:** consensus repeat of all arrays in the group (might be the reverse complement from the one shown in CRISPRCasdb)

**array_names:** list of names of all arrays in the group, structure: "CRISPR-ID_start-pos_end-pos" as reported by CRISPRCasdb for individual CRISPR arrays

**nb_of_arrays:** number of arrays in group

**cas_type:** Cas type assigned according to closest (< 10000 bp distance to array) Cas locus.

**kingdom:** kingdom of samples in group

**genus:** genus of samples in group

**array_species:** list of species reported in CRISPRCasdb for each array

**forward_ln_lh_bdm_–_reversed_ln_lh_bdm:** value of test statistic used by *CRISPR-evOr* for orientation prediction (positive indicates forward orientation, negative indicates reversed orientation)

**crispr-evOr_prediction:** prediction by *CRISPR-evOr* using the test statistic (threshold for confident prediction of $c = 5$)

**group_pam_prediction:** assigned PAM-orientation for whole group (based on individual predictions as described in the supplement), "not available" means that there were no samples in this group which are also in the PAM-dataset

**group_direction_prediction:** assigned orientation by CRISPRDirection for the whole group (based on individual predictions as described in the supplement)

**group_strand_prediction:** assigned orientation by CRISPRstrand for the whole group (based on individual predictions as described in the supplement)

**group_strand_confidence:** CRISPRstrand assigns a confidence value for its prediction. Here shown is the one assigned for the whole group (CRISPRstrand's confidence is always consistent within groups)

**array_toolname_prediction:** all methods excluding *CRISPR-evOr* are single-array-based, list of dictionaries with orientation predictions by all single-array-based methods for each array in "array_names" (same order); for "pam-prediction": "not available" means that the array is was not found in the PAM-dataset; CRISPRstrand has additional individual confidence values

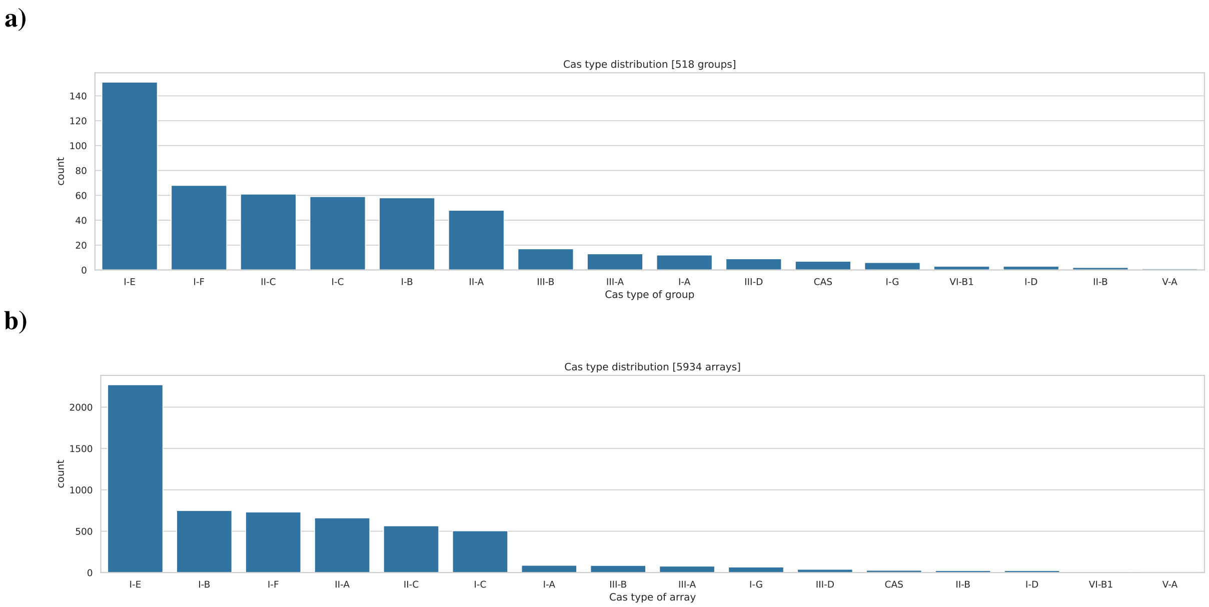


**Supplementary Fig. S1.** We show the count of groups (**a)**) and spacer arrays (**b)**) according to their Cas type of the considered CRISPRCasdb dataset. Note that groups are composed of arrays of the same Cas type.

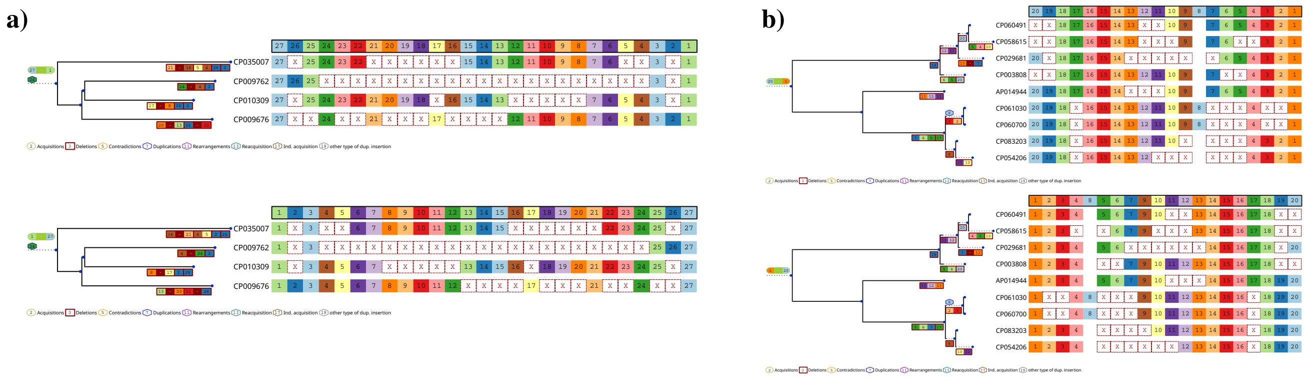


**Supplementary Fig. S2.** We show examples with likelihood ratio $L_{\text{ratio}} = 0$ on core-genome trees with the forward (above) and reverse (below) reconstructions. To make the figures more concise, acquisition and deletion events in the tree are consolidated, e.g. 27 - 1 in green (red) indicates that spacers 27 through 1 were acquired (deleted) at this branch. In **a)**, the first and last spacer are present in every array and thus force the reconstruction to be the same in forward and reverse orientation. In **b)**, although the first/last spacer are not present in all arrays, SpacerPlacer finds the most likely reconstruction to be with spacer 20 in the root (due to CP029681), leading to a similar outcome to a). The shown examples are groups of a) *Staphylococcus schleiferi* (Cas type II-C) and b) a mixture of *Staphylococcus aureus*, *Staphylococcus pseudintermedius* and *Staphylococcus schleiferi* (Cas type III-A)

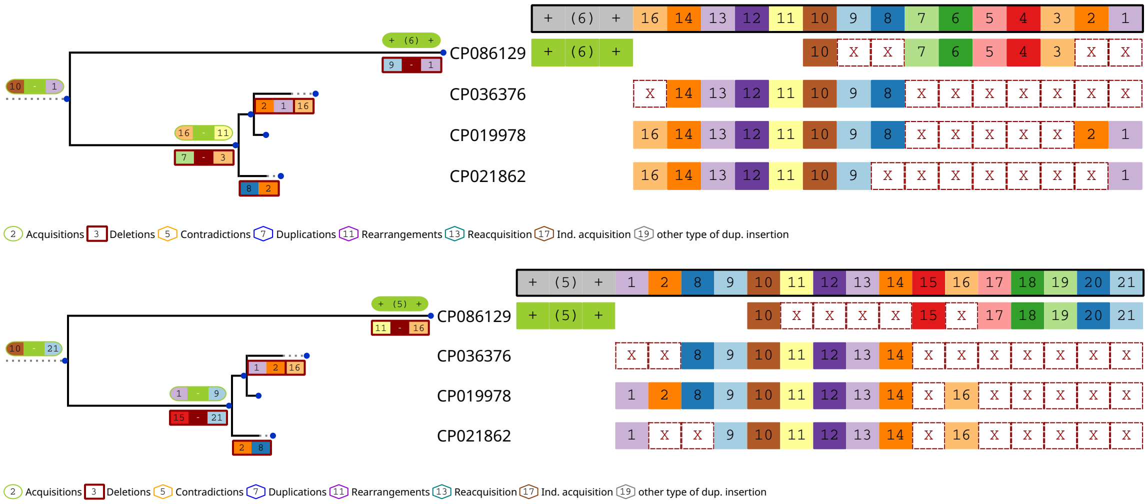


**Supplementary Fig. S3. Example of a group with low confidence orientation prediction.** We show an example where *CRISPR-evOr* is not confident. Both orientations lead to reasonable reconstructions and although *CRISPR-evOr* is able to give a suggestion ($|L_{\text{ratio}}| = 1.50$ in favor of the lower reconstruction), it will not be confident according to our confidence threshold. To make the figures more concise, acquisition and deletion events in the tree are consolidated, e.g. 10 - 1 in green (red) indicates that spacers 10 through 1 were acquired (deleted) at this branch. At the leafs and in the alignment "+ (x) +" indicates that x spacers were acquired (which are not found in any other array), e.g. in the top and bottom figure (6) and (5) represent spacers 15, 17 - 21 and 3 - 7 respectively. The shown example is a group of *Streptococcus agalactiae* (Cas type I-C) from CRISPRCasdb.

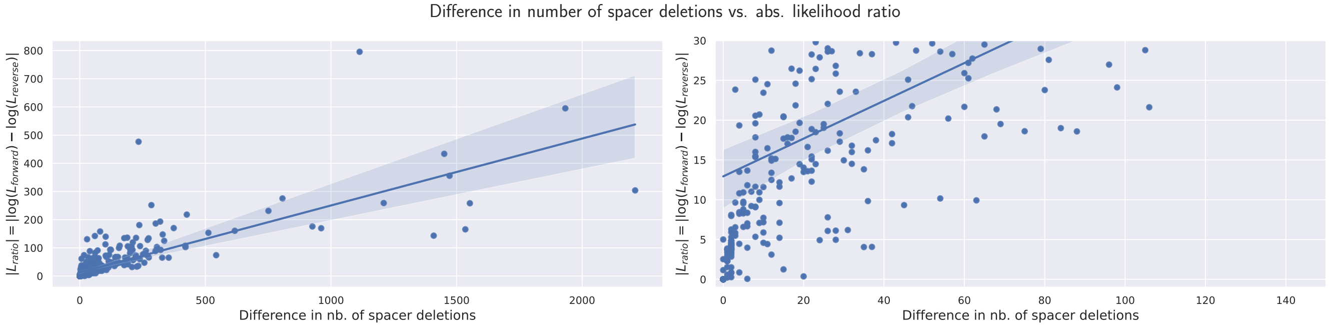


**Supplementary Fig. S4.** We show the difference of spacer deletions between the forward and reverse reconstruction versus the absolute value of the respective difference in likelihood ratios $|L_{\text{ratio}}|$ for the CRISPRCasdb dataset. Each scatter point indicates the respective values for one group in the CRISPRCasdb dataset. Furthermore, we give a linear regression with shaded 95% confidence interval based on 1000-fold bootstrapping. The left side shows all of the dataset and the right side is zoomed in for better readability. No point with more than 150 difference in spacer deletions has a absolute likelihood ratio less than 30. Note, that the likelihood ratio increases rapidly with increasing difference in the number of spacer deletions and after $|L_{\text{ratio}}| > 5$ most decisions are made based on more than 5 spacer deletions.

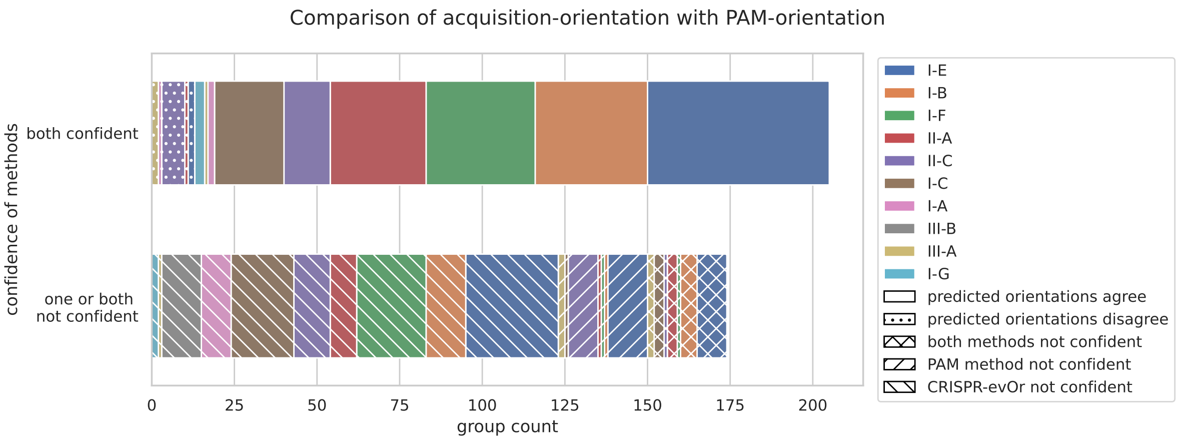


**Supplementary Fig. S5.** Complementary to Figure 3 in the main manuscript, we show the performance and confidence comparison between *CRISPR-evOr* and PAM-orientation in terms of groups. Note, that the PAM-orientation is assigned to groups by taking the confident predictions of arrays within the group. There are no contradictions of PAM-orientations within groups, but the amount of groups with high confidence PAM-orientation predictions is somewhat inflated since the group prediction can rely on only very few arrays with confident predictions. See Supplementary Note 1C for remarks about how we arrive at the group orientation.

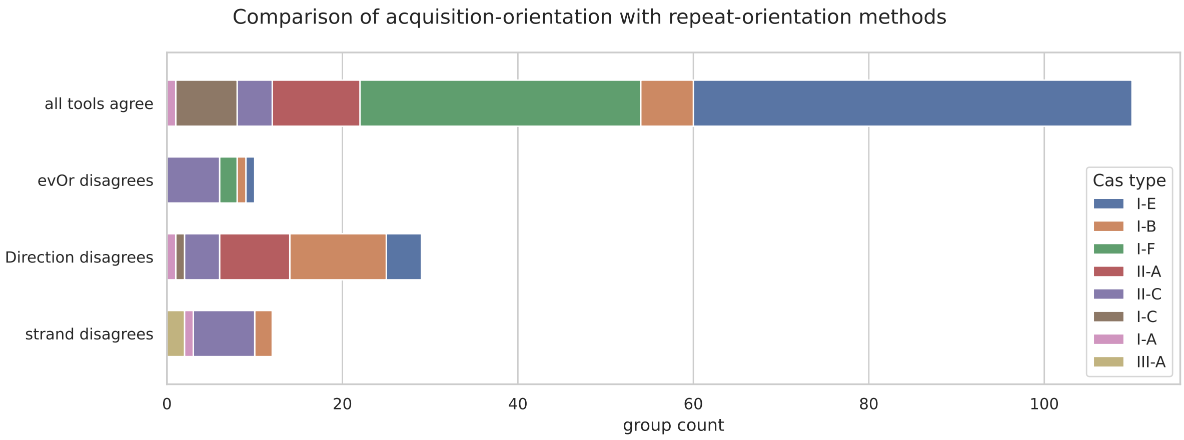


**Supplementary Fig. S6.** Complementary to Figure 4 in the main manuscript, we show the performance comparison between CRISPRDirection, CRISPRstrand and *CRISPR-evOr* in terms of groups. We only consider groups where all three tools have confident predictions. Note, that the repeat-orientations are assigned to groups by taking the confident predictions of arrays within the group. There are no contradictions of repeat-orientations within groups, but the amount of groups with high confidence repeat-orientation predictions is somewhat inflated since the group prediction can rely on only very few arrays with confident predictions. See Supplementary Note 1C for remarks about how we arrive at the group orientation.

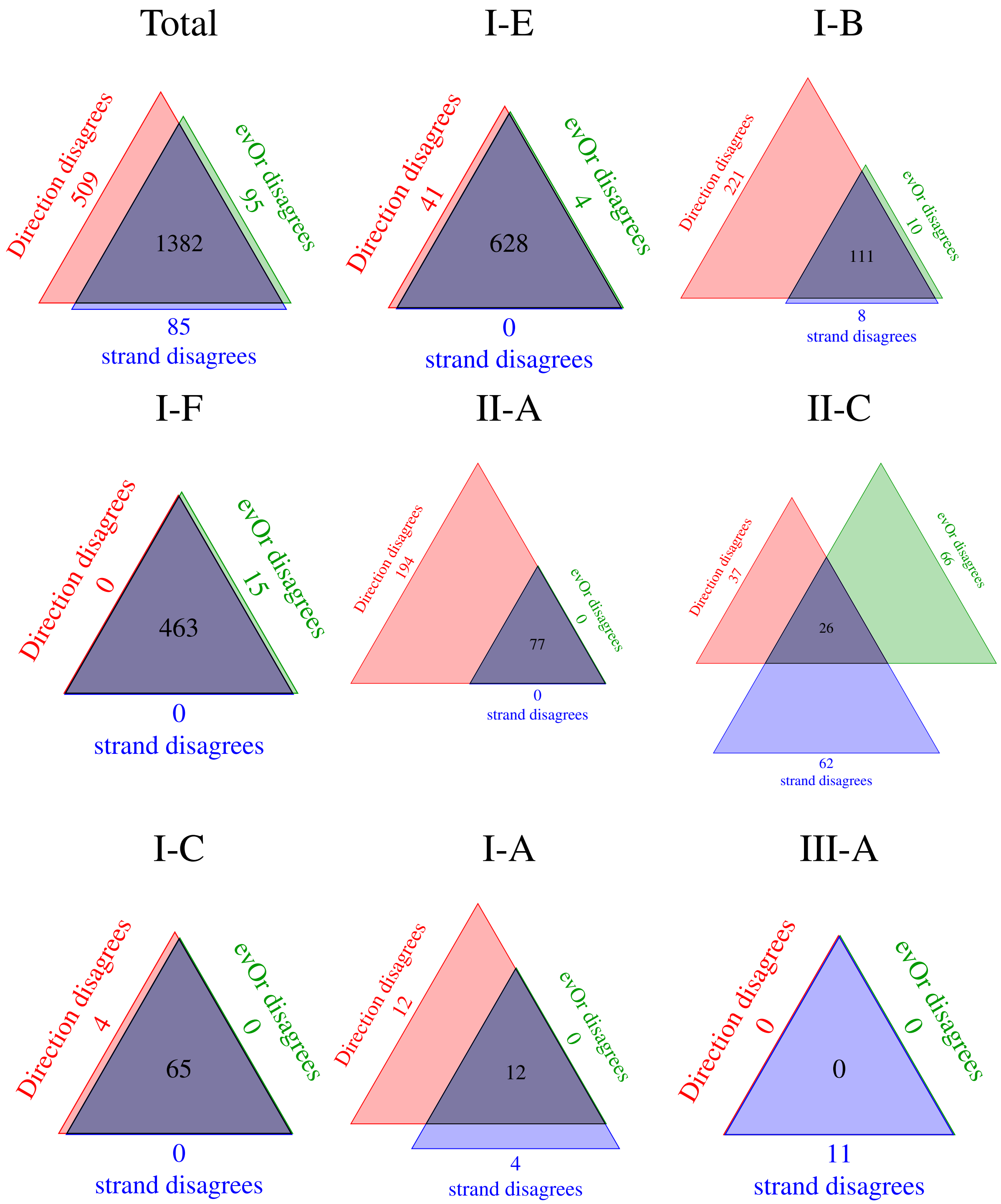


**Supplementary Fig. S7.** This Figure is equivalent to Figure 4 in the main manuscript. We show (as Venn diagrams) the agreement of orientation predictions between CRISPRDirection, CRISPRstrand and *CRISPR-evOr* where all methods have high confidence broken down according to Cas type. The Venn diagrams clearly show substantial differences in agreement for different Cas types. For type I-E and I-F the tools mostly agree. However, CRISPRDirection disagrees significantly for I-B, II-A and all tools quite often disagree for II-C. There are indications that CRISPRstrand may not accurately predict the orientation of type III-A arrays; however, the sample size is very small.

**a)**

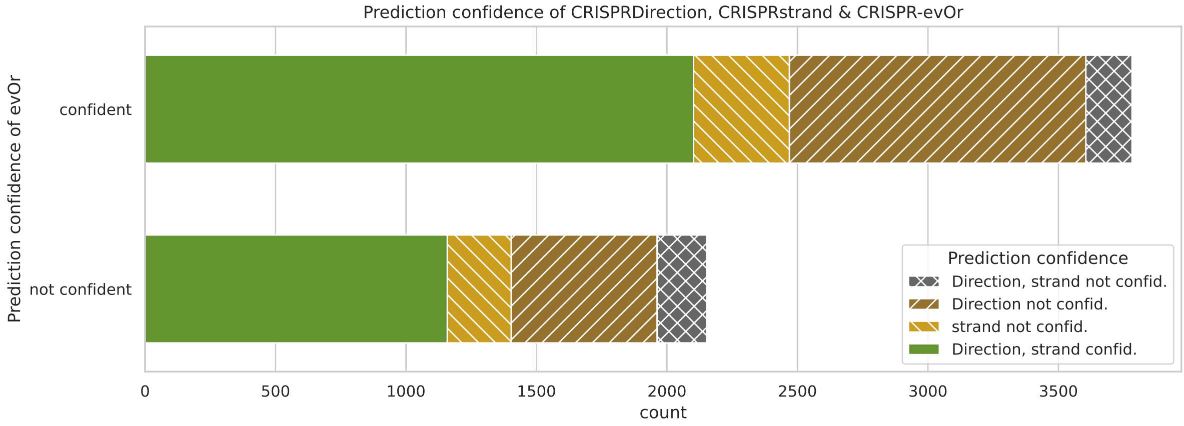


**b)**

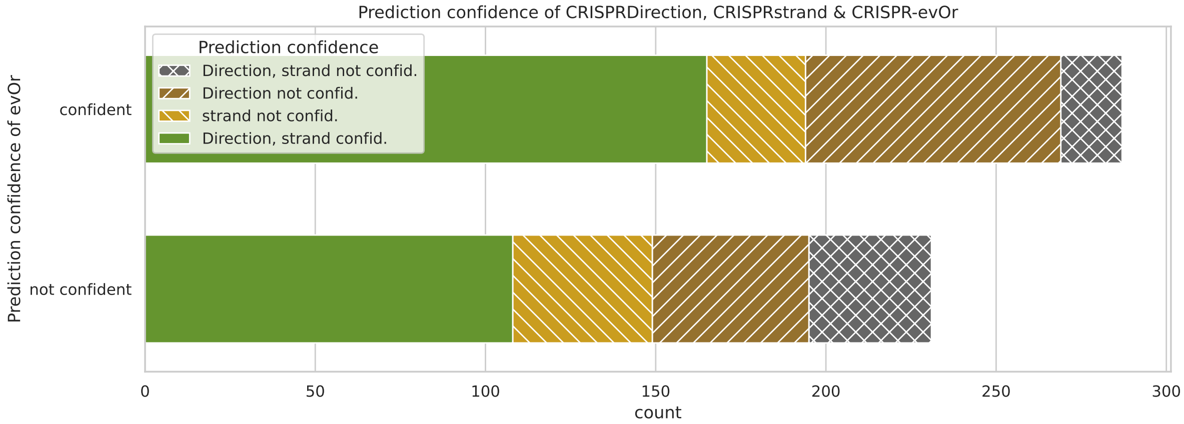


**Supplementary Fig. S8. Confidence comparison between *CRISPR-evOr* and repeat based methods.** We show a breakdown of arrays (**a)**) and groups (**b)**) in the CRISPRCasdb dataset where *CRISPR-evOr*, CRISPRDirection, and CRISPRstrand are able to make confident decisions. It shows that there is a large extent of the dataset where *CRISPR-evOr* is confident in its prediction. Furthermore, there is a substantial amount of data, where at least one of the other methods is not decisive.

a)

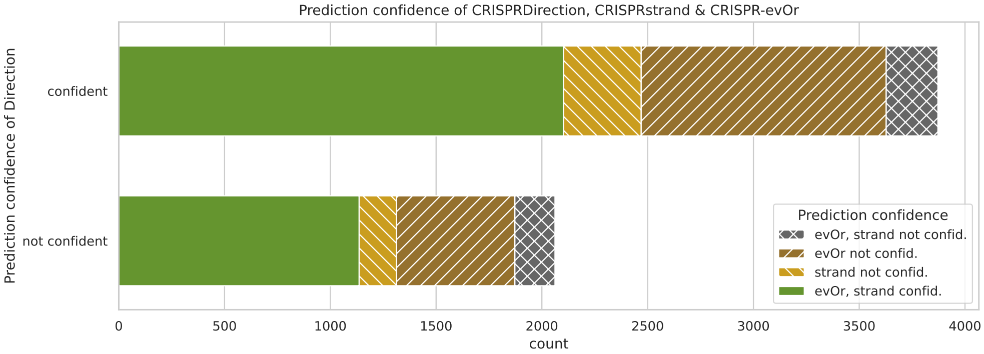


b)

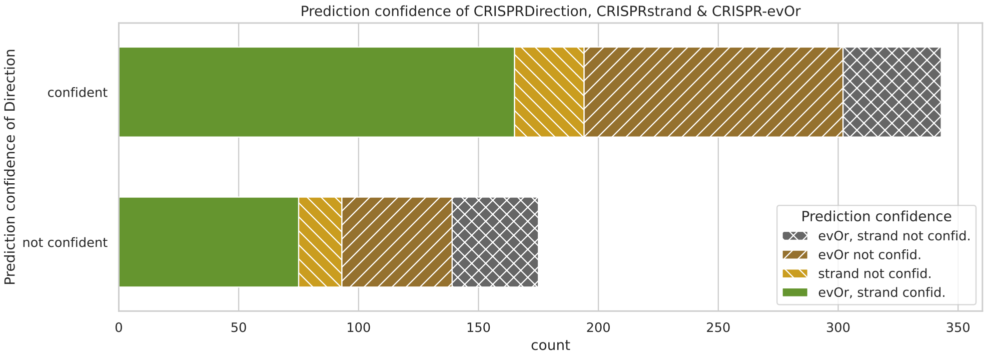


c)

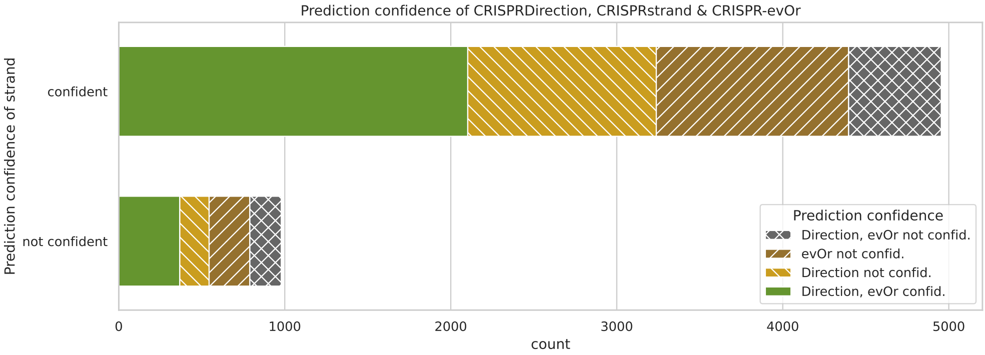


d)

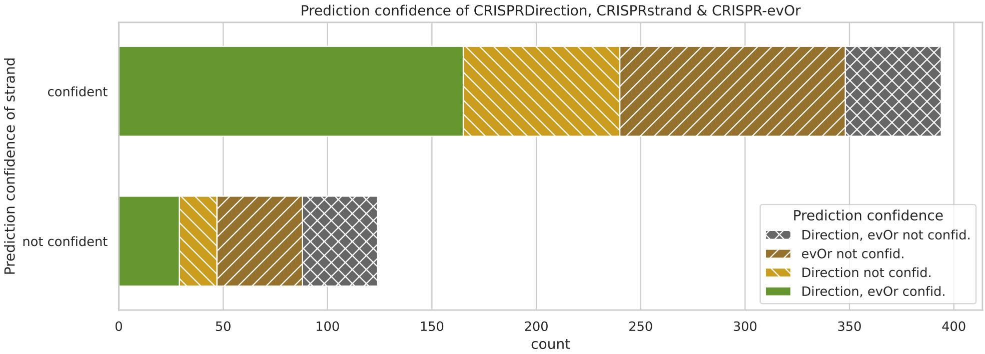


**Supplementary Fig. S9. Confidence comparison between *CRISPR-evOr* and repeat based methods (other y-axis).** We show a breakdown of arrays (**a)**) and groups (**b)**) in the CRISPRCasdb dataset where *CRISPR-evOr*, CRISPRDirection, and CRISPRstrand are able to make confident decisions with CRISPRDirection on the y-axis. Below we show the same statistics (arrays **c)** and groups (**d)**) with CRISPRstrand on the y-axis.

**a)**

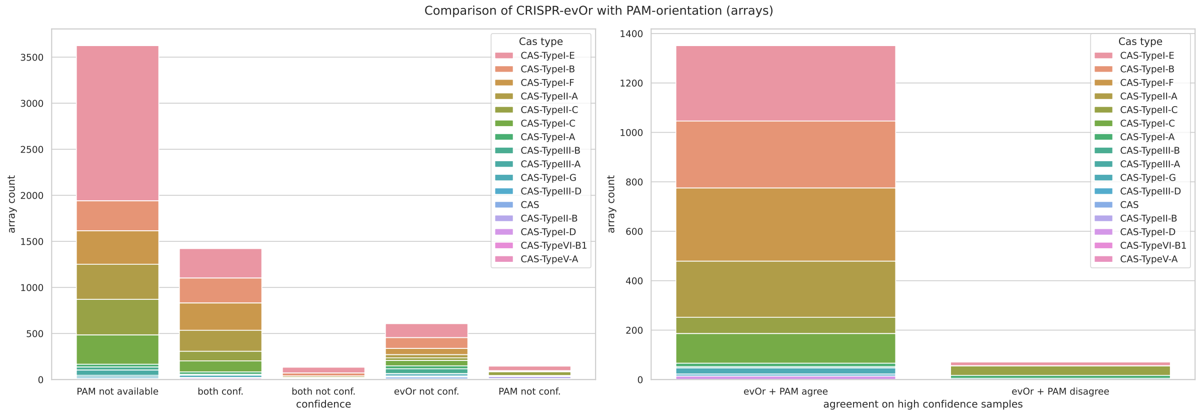


**b)**

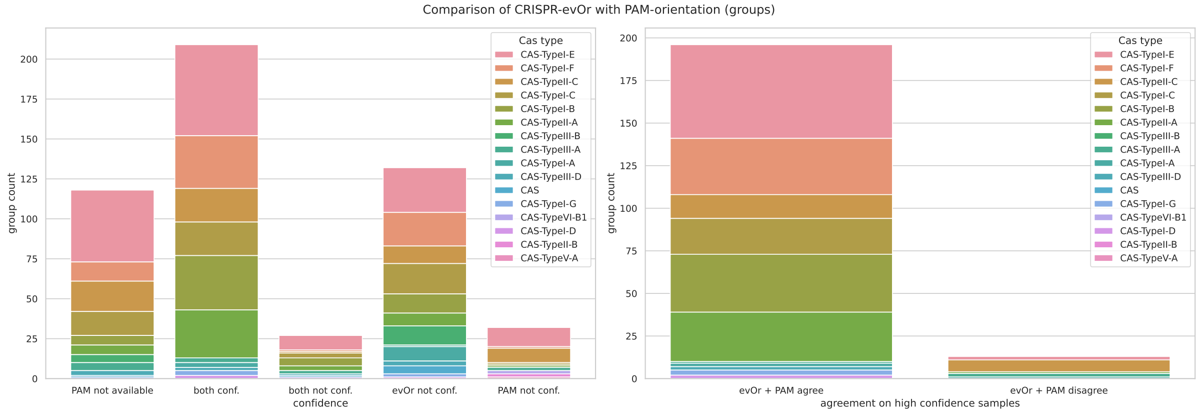


**Supplementary Fig. S10.** Comparison between acquisition-orientation and PAM-orientation broken down according to Cas types on our CRISPRCasdb dataset. On the left we show the confidence on the dataset and on the right the agreement of the predictions for high confidence samples. Count in **a)** arrays and **b)** groups. Additionally to the figure in the main manuscript we show the Cas types with few samples and the amount of arrays/groups for which no prediction is available ("PAM not available") in the PAM-orientation dataset [2].

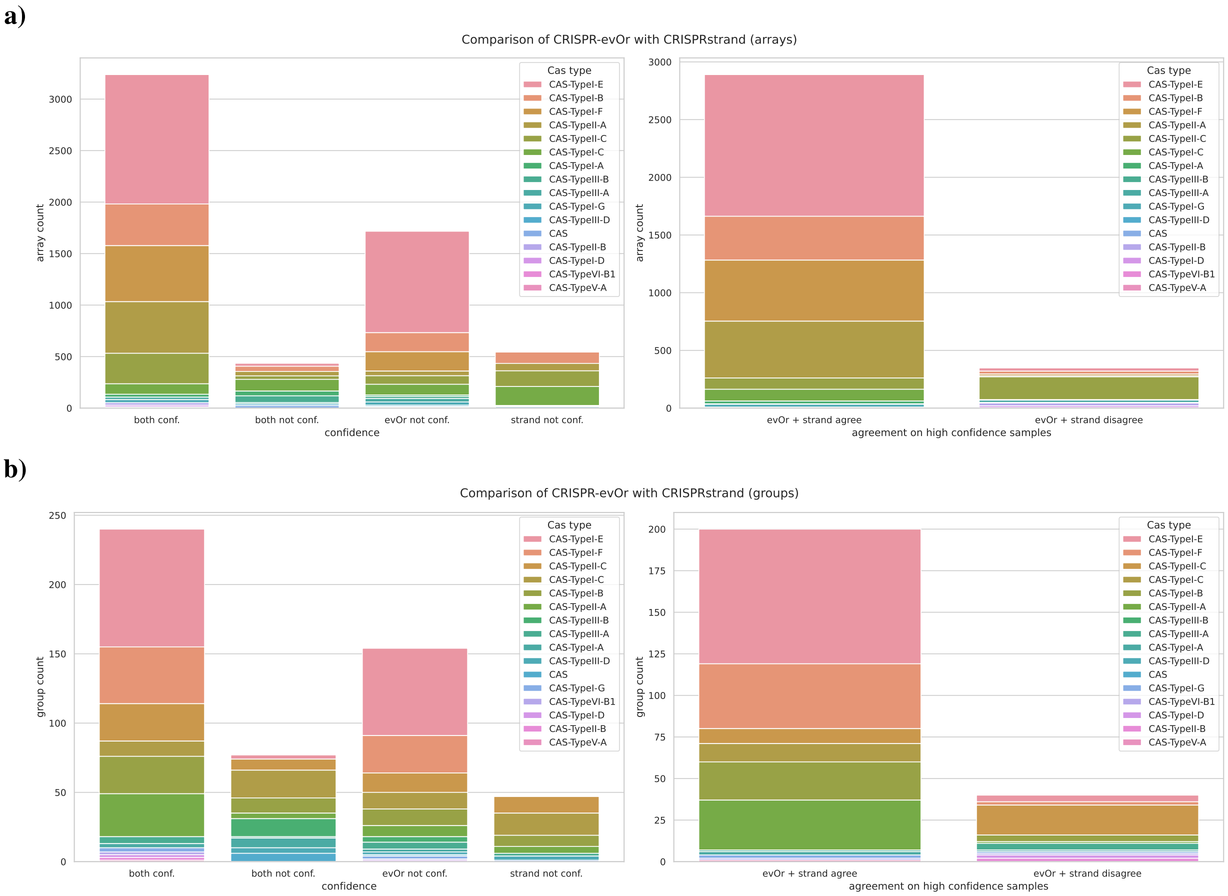


**Supplementary Fig. S11.** Comparison between acquisition-orientation and CRISPRstrand broken down according to Cas types on our CRISPRCasdb dataset. On the left we show the confidence on the dataset and on the right the agreement of the predictions for high confidence samples. Count in **a)** arrays and **b)** groups. Additionally to the figure in the main manuscript we show the Cas types with few samples.

**a)**

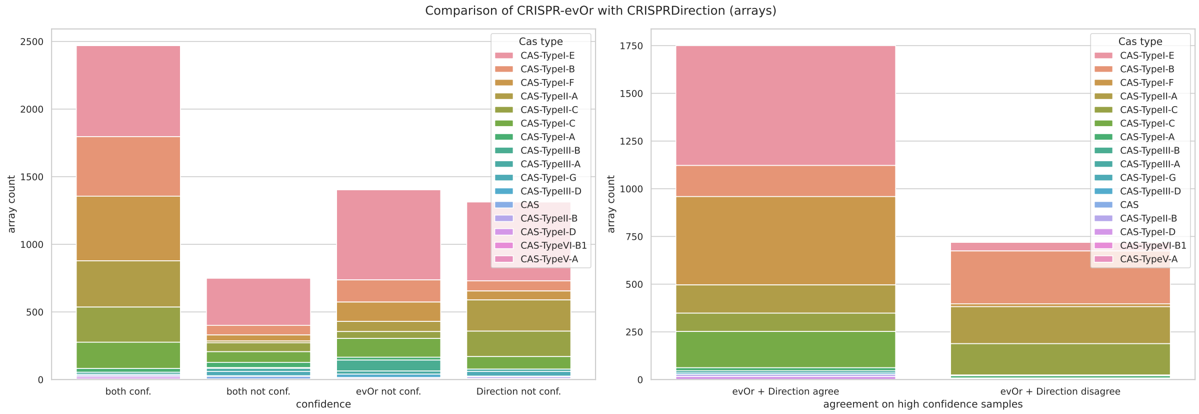


**b)**

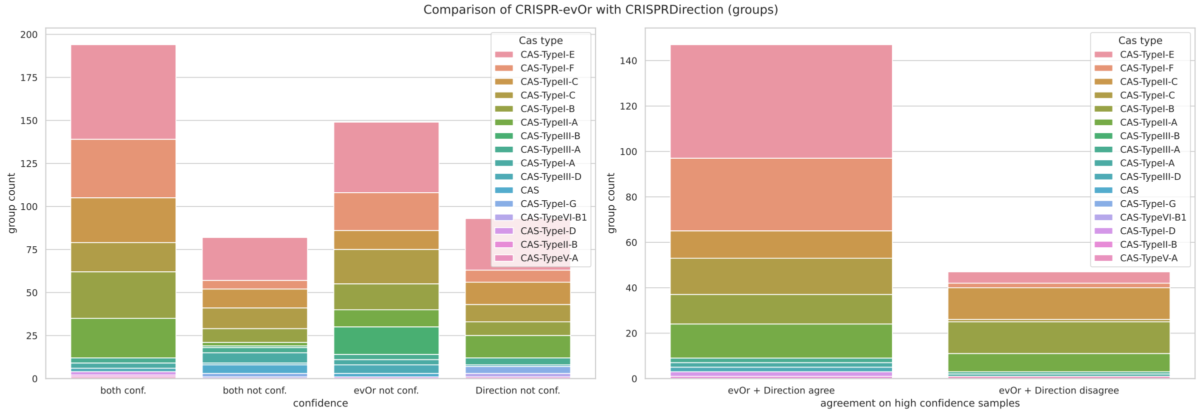


**Supplementary Fig. S12.** Comparison between acquisition-orientation and CRISPRDirection broken down according to Cas types on our CRISPRCasdb dataset. On the left we show the confidence on the dataset and on the right the agreement of the predictions for high confidence samples. Count in **a)** arrays and **b)** groups. Additionally to the figure in the main manuscript we show the Cas types with few samples.

**a)**

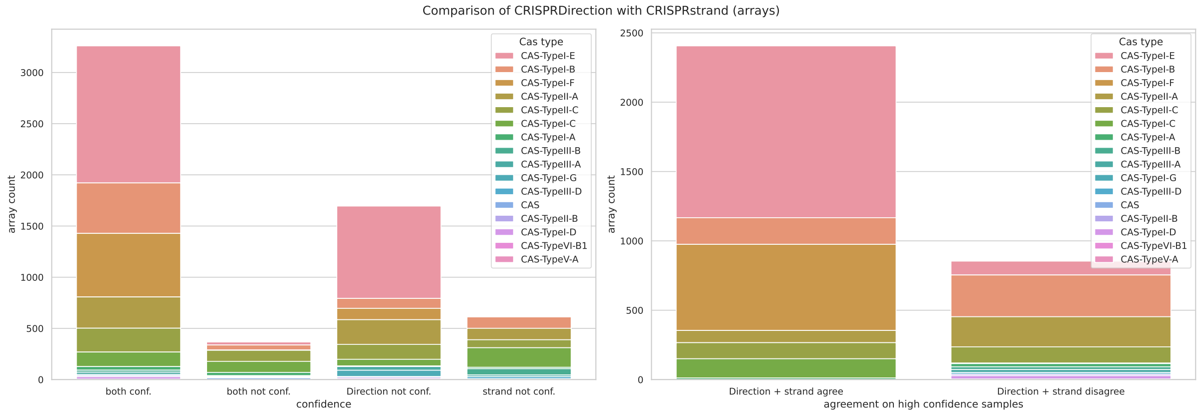


**b)**

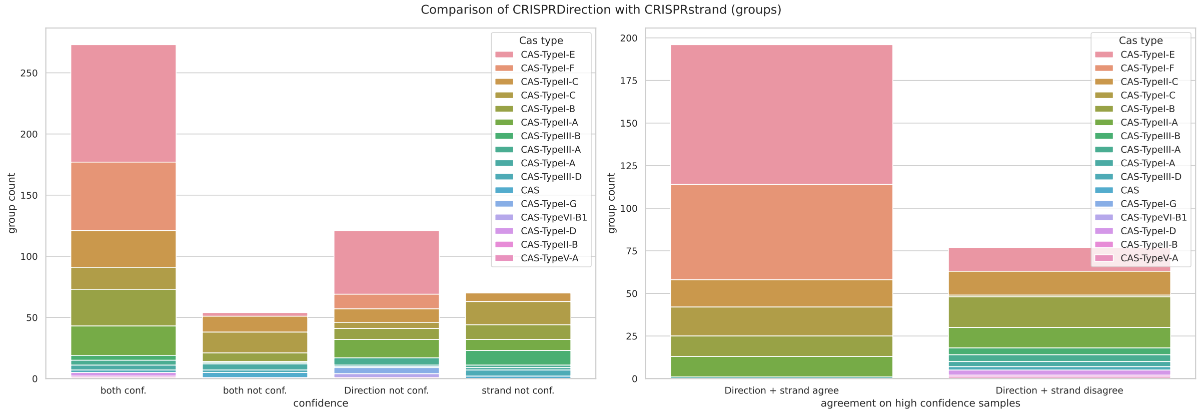


**Supplementary Fig. S13.** Comparison between CRISPRDirection and CRISPRstrand broken down according to Cas types on our CRISPRCasdb dataset. On the left we show the confidence on the dataset and on the right the agreement of the predictions for high confidence samples. Count in **a)** arrays and **b)** groups. Additionally to the figure in the main manuscript we show the Cas types with few samples.

**a)**

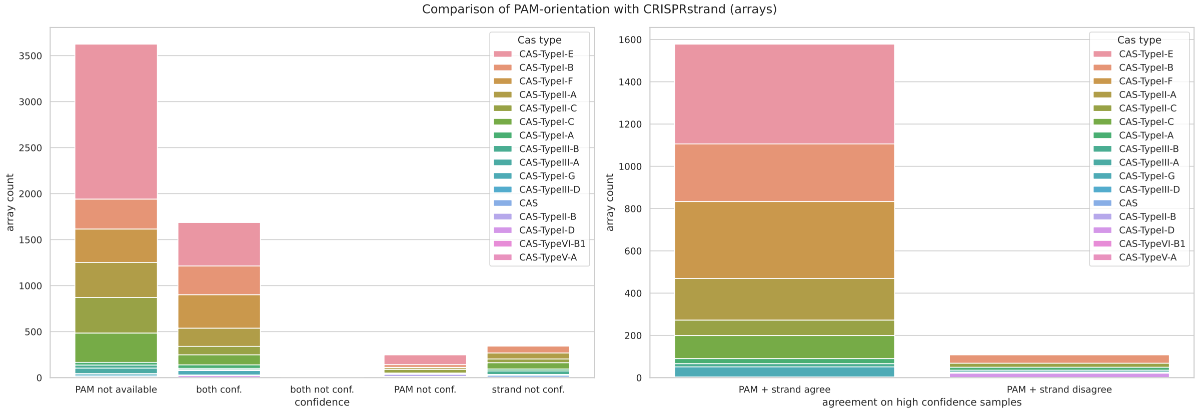


**b)**

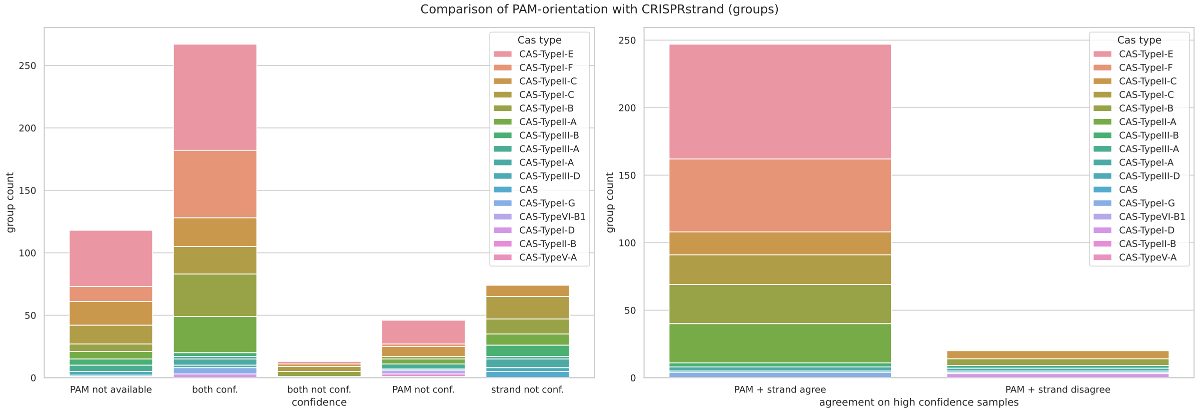


**Supplementary Fig. S14.** Comparison between PAM-orientation and CRISPRstrand broken down according to Cas types on our CRISPRCasdb dataset. On the left we show the confidence on the dataset and on the right the agreement of the predictions for high confidence samples. Count in **a)** arrays and **b)** groups. Additionally to the figure in the main manuscript we show the Cas types with few samples and the amount of arrays/groups for which no prediction is available (“PAM not available”) in the PAM-orientation dataset [2]. Note, that the amount of arrays where both tools are not confident is so small (32), that it might not be visible.

**a)**

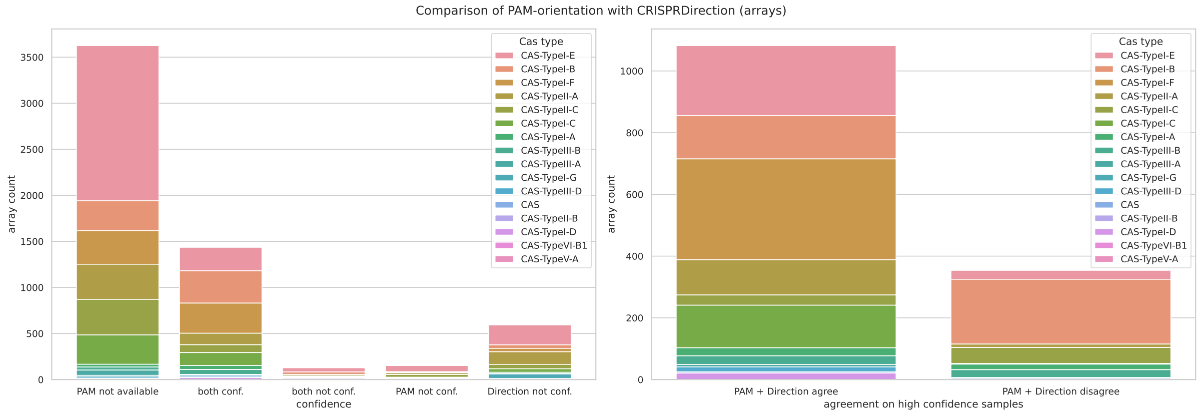


**b)**

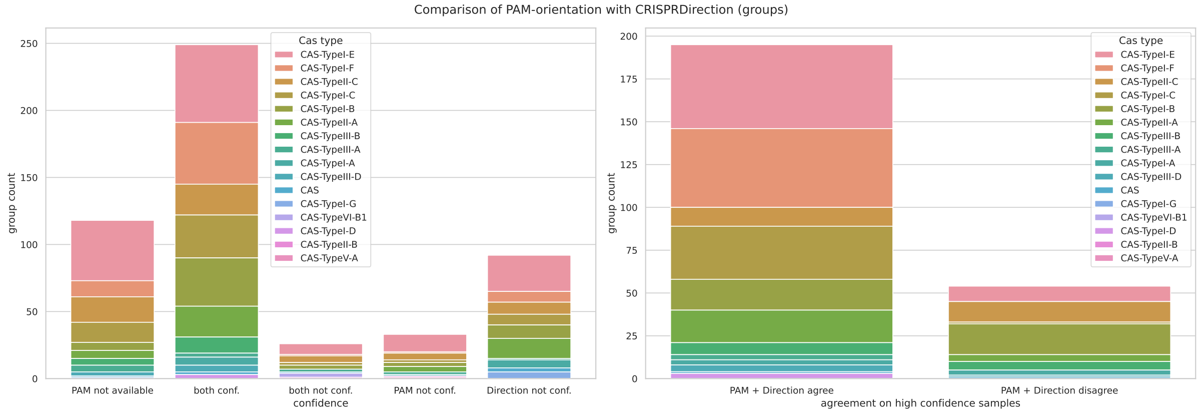


**Supplementary Fig. S15.** Comparison between PAM-orientation and CRISPRDirection broken down according to Cas types on our CRISPRCasdb dataset. On the left we show the confidence on the dataset and on the right the agreement of the predictions for high confidence samples. Count in **a)** arrays and **b)** groups. Additionally to the figure in the main manuscript we show the Cas types with few samples and the amount of arrays/groups for which no prediction is available ("PAM not available") in the PAM-orientation dataset [2].

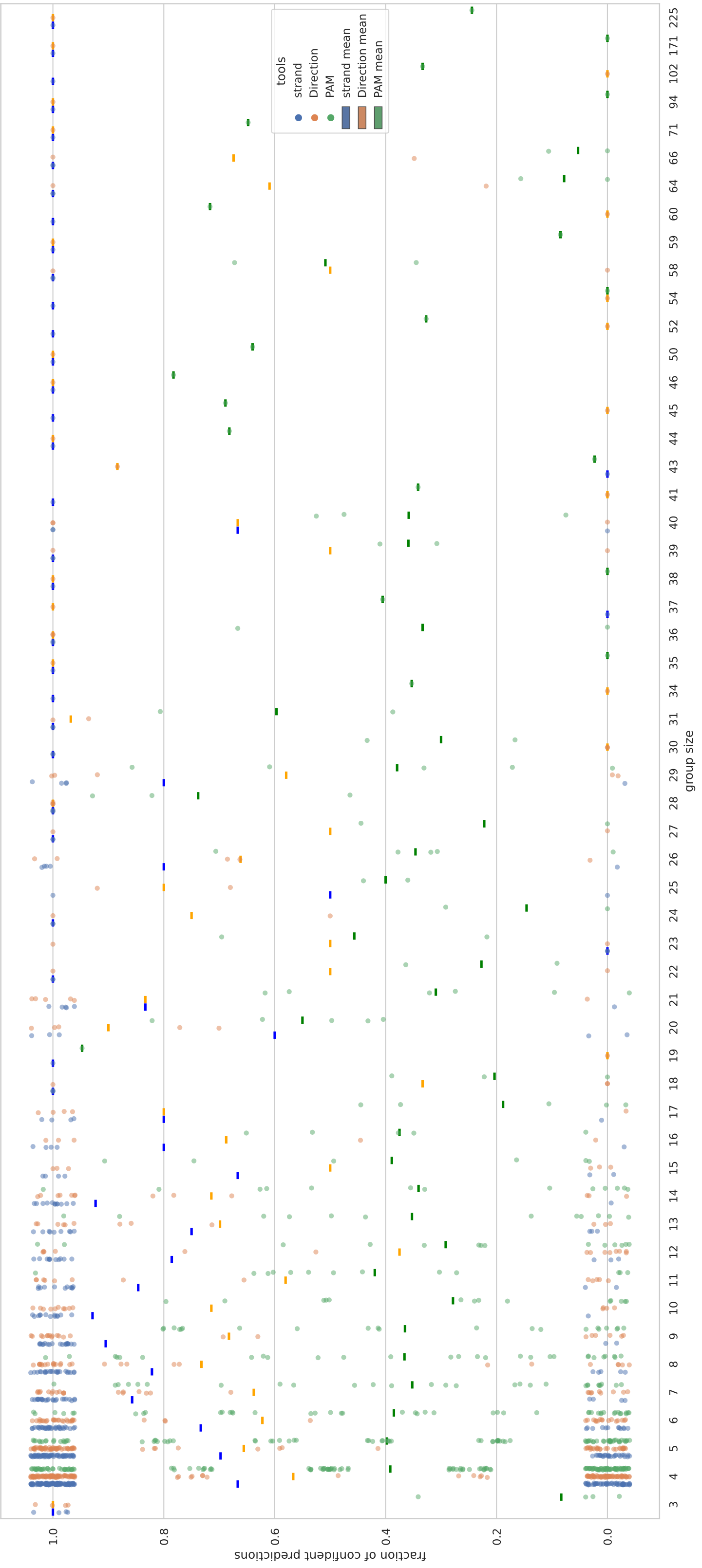


**Supplementary Fig. S16.** We show the fraction of confident predictions for each group and array-based tool versus the number of arrays in the respective group on the complete CRISPRCasdb dataset. Moreover, we show the means of the fraction of confident predictions for each group size and tool as the respective colored dashes. Note that we introduced some noise to the individual group percentages to make individual groups distinguishable, i.e. the dots y-value is the respective closest multiple of $k/n$, where $n$ is the group size and $k \in \{0, \ldots n\}$. For PAM-orientation, both the uncertain predictions and the arrays for which no prediction was available from the dataset of Vink et al. are labeled as uncertain and thus contribute to the fractions.

### Bibliography


1. Axel Fehrenbach, Alexander Mitrofanov, Omer S Alkhnbashi, Rolf Backofen, and Franz Baumdicker. SpacerPlacer: ancestral reconstruction of CRISPR arrays reveals the evolutionary dynamics of spacer deletions. *Nucleic Acids Research*, page gkae772, September 2024. ISSN 0305-1048, 1362-4962. doi: 10.1093/nar/gkae772.
2. Jochem N. A. Vink, Jan H. L. Baijens, and Stan J. J. Brouns. PAM-repeat associations and spacer selection preferences in single and co-occurring CRISPR-Cas systems. *Genome Biology*, 22(1):281, December 2021. ISSN 1474-760X. doi: 10.1186/s13059-021-02495-9.
